# FinisherSC : A repeat-aware tool for upgrading de-novo assembly using long reads

**DOI:** 10.1101/010215

**Authors:** Ka-Kit Lam, Kurt LaButti, Asif Khalak, David Tse

## Abstract

We introduce FinisherSC, a repeat-aware and scalable tool for upgrading de-novo assembly using long reads. Experiments with real data suggest that FinisherSC can provide longer and higher quality contigs than existing tools while maintaining high concordance.

**Availability:** The tool and data are available and will be maintained at http://kakitone.github.io/finishingTool/

## 1 INTRODUCTION

In de-novo assembly pipelines for long reads, reads are often trimmed or thrown away. Moreover, there is no evidence that state-of-the-art assembly pipelines are data-efficient. In this work, we ask whether state-of-the-art assembly pipelines for long reads have already used up all the available information from raw reads to construct assembly of the highest possible quality. To answer this question, we first collect output contigs from the HGAP (Chin et al., 2013) pipeline and the associated raw reads. Then, we pass them into our tool FinisherSC (Lam et al., 2014b) to see if higher quality assemblies can be consistently obtained after post-processing.

## 2 METHODS

### 2.1 Usage and pipeline

FinisherSC is designed to upgrade de-novo assembly using long reads (e.g. PacBio^®^ reads). It is especially suitable for data consisting of a single long reads library. Input to FinisherSC are contigs (contigs.fasta) constructed by an assembler and all the raw reads with adaptors removed (raw_reads.fasta). Output of FinisherSC are upgraded contigs (improved3.fasta) which are expected to be of higher quality than its input (e.g. longer N50, longer longest contigs, fewer number of contigs, high percentage match with reference, high genome fraction, etc). In Fig 1, we show an example pipeline in which FinisherSC can fit. As shown in Fig 1, FinisherSC can be readily incorporated into state-of-the-art assembly pipelines (e.g. PacBio^®^HGAP).

**Fig. 1.**
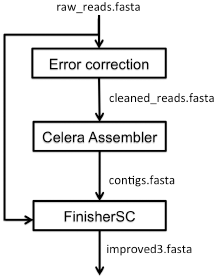
Pipeline where FinisherSC can fit in

### 2.2 Algorithm and features

The algorithm of FinisherSC is summarized in Alg 1. Detailed description of the algorithm is in the supplementary materials. We summarize the key features of FinisherSC as follows.

- Repeat-aware: FinisherSC uses a repeat-aware rule to define overlap. It uses string graphs to capture overlap information and to handle repeats so that FinisherSC can robustly merge contigs. There is an optional component, X-phaser (Lam et al., 2014a), that can resolve long approximate repeats with two copies by using the polymorphisms between them. There is also an optional component, T-solver, that can resolve tandem repeat by using the copy count information.
- Data-efficient: FinisherSC utilizes all the raw reads to perform re-layout. This can fill gaps and improve robustness in handling repeats.
- Scalable: FinisherSC streams raw reads to identify relevant reads for re-layout and refined analysis. MUMMER (Kurtz et al., 2004) does the core of the sequence alignment. Although MUMMER is single threaded, we provide an option to segment the files and run multiple MUMMER jobs in parallel. These techniques allow FinisherSC to be easily scalable to high volume of data.

#### Algorithm 1: Main flow of FinisherSC

**Input:** contigs.fasta, raw_reads.fasta

**Output:** improved3.fasta

1. Filter completely embedded contigs
2. Form a string graph with the BEST successors/predecessors as edges
3. Condense the string graph by contracting edges with both in-degree and out-degree being 1
4. Use raw reads to declare potential successors/predecessors of dangling contigs
5. Merge contigs (with gaps filled by reads) when they respectively only have 1 successor/1 predecessor
6. Form a string graph with ALL successors/predecessors as edges
7. Merge contigs with only 1 predecessor or 1 successor and each has no more than two competing edges

## 3 RESULTS AND DISCUSSION

### 3.1 Experimental evaluation on bacterial genomes

We evaluated the performance of FinisherSC as follows. Raw reads were processed according to the pipeline in Fig 1. They were first error corrected and then assembled into contigs by an existing pipeline (i.e. HGAP). Contigs were upgraded using FinisherSC and evaluated for quality with Quast (Gurevich et al., 2013). The data used for assessment are real PacBio^®^ reads. These include data recently produced at JGI and data available online supporting the HGAP publication. We compared the assembly quality of the contigs coming out from the Celera assembler (Myers et al., 2000) of HGAP pipeline, the upgraded contigs by FinisherSC and the upgraded contigs by PBJelly (English et al., 2012). A summary of the evaluation is shown in Fig 2. More details can be found in the supplementary materials. We find that FinisherSC can upgrade the assembly from HGAP without sacrifice on accuracy on these test cases. Moreover, the upgraded contigs by FinisherSC are generally of higher quality than those upgraded by PBJelly. This suggests that there is extra information from the reads that is not fully utilized by state-of-the-art assembly pipelines for long reads.

**Fig. 2.**
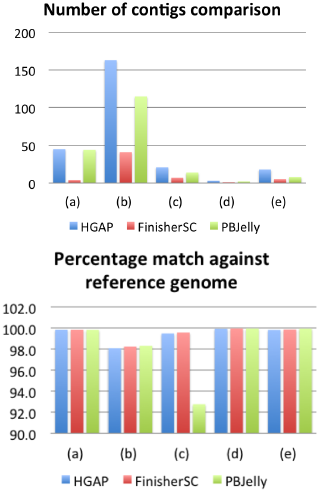
Experimental evaluation on bacterial genomes. (a,b) : Pedobacter heparinus DSM 2366 (PacBio^®^ long reads from JGI) (c, d, e) : Escherichia coli MG 1655, Meiothermus ruber DSM 1279, Pedobacter heparinus DSM 2366 (PacBio^®^ long reads supporting the HGAP publication).

### 3.2 Experiments on scalability

We tested the scalability of FinisherSC by applying it to handle larger genomes. The data used are the benchmark data available on PacBio Devnet. We run FinisherSC with the option of using 20 threads (-par 20) on a server computer. The server computer is equipped with 64 cores of CPU at clock rate of 2.4-3.3GHz and 512GB of RAM. The running time is tabulated in Table 1.

**Table 1.** Summary of running time for the experiments on scalability

| Genome name | Genome size (Mbp) | Size of reads (Gbp) | Running time (hours) |
| --- | --- | --- | --- |
| Caenorhabditis elegans | 104 | 7.65 | 23 |
| Drosophila | 138 | 2.27 | 9.4 |
| Saccharomyces cerevisiae | 12.4 | 1.40 | 0.66 |

### 3.3 Discussion

Although FinisherSC was originally designed to improve de-novo assembly by long reads, it can also be used to scaffold long contigs (formed by short reads) using long reads. For that use case, we note that the contigs formed by short reads can sometimes have length shorter than the average length of long reads. Therefore, we suggest users to filter out those short contigs before passing them into FinisherSC.

## 4 SUPPLEMENTARY MATERIALS

### 4.1 Summary of dataflow in FinisherSC and key experimental results

We visualize in Figure 3 the flow of FinisherSC. Moreover, we summarize the key experimental results in Table 2.

**Fig. 3.**
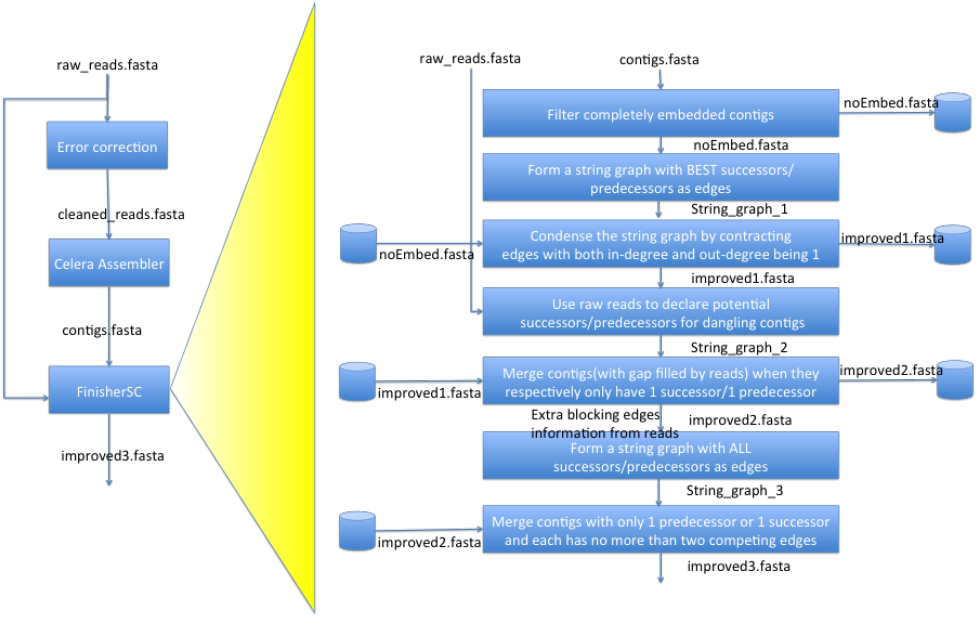
Summary of data flow of FinisherSC

**Table 2.** Experimental evaluation results. (a,b) : Pedobacter heparinus DSM 2366 (recent real long reads from JGI) (c, d, e) : Escherichia coli MG 1655, Meiothermus ruber DSM 1279, Pedobacter heparinus DSM 2366 (real long reads supporting the HGAP publication). Detailed analysis by Quast is shown in the supplementary material.

| Genome name | (a) | (b) | (c) | (d) | (e) |
| --- | --- | --- | --- | --- | --- |
| Genome size | 5167383 | 5167383 | 4639221 | 3097457 | 5167383 |
| Coverage of raw reads | 50 | 30 | 50 | 55 | 53.1 |
| Coverage of corrected reads | 32.93 | 17.74 | 10.1 | 11.3 | 9.5 |
| Coverage of input to Celera | 32.93 | 17.74 | 10.1 | 11.3 | 9.5 |
| N50 of HGAP output | 4097401 | 89239 | 392114 | 1053479 | 1403814 |
| N50 of FinisherSC upgraded output | 5168551 | 215810 | 1525398 | 3099349 | 2913716 |
| N50 of PBJelly upgraded output | 4099674 | 145441 | 1200847 | 1715191 | 3343452 |
| Number of contigs of HGAP output | 45 | 163 | 21 | 3 | 18 |
| Number of contigs of FinisherSC upgraded output | 4 | 41 | 7 | 1 | 5 |
| Number of contigs of PBJelly upgraded output | 44 | 115 | 14 | 2 | 8 |
| Length of the longest contig of HGAP output | 4097401 | 254277 | 1241016 | 1390744 | 2103385 |
| Length of the longest contig of FinisherSC upgraded output | 5168551 | 637485 | 2044060 | 3099349 | 2913716 |
| Length of the longest contig of PBJelly upgraded output | 4099674 | 495596 | 1958341 | 1715191 | 3343452 |
| Percentage match of HGAP output against reference | 99.87 | 98.11 | 99.51 | 99.96 | 99.85 |
| Percentage match of FinisherSC upgraded output against reference | 99.87 | 98.27 | 99.60 | 99.99 | 99.89 |
| Percentage match of PBJelly upgraded output against reference | 99.86 | 98.34 | 92.77 | 99.98 | 99.97 |
| Total length of HGAP output | 5340498 | 5536634 | 4689701 | 3102769 | 5184825 |
| Total length of FinisherSC upgraded output | 5212355 | 5139491 | 4660679 | 3099349 | 5167414 |
| Total length of PBJelly upgraded output | 5383836 | 5821106 | 4718818 | 3106774 | 5210862 |

**Table 3.** (a) in Table 2. All statistics are based on contigs of size ≥ 500 bp, unless otherwise noted (e.g., ”# contigs (≥ 0 bp)” and ”Total length (≥ 0 bp)” include all contigs).

| Assembly | HGAP | FinisherSC | PBJelly |
| --- | --- | --- | --- |
| # contigs ($\geq 0$ bp) | 45 | **4** | 44 |
| # contigs ($\geq 1000$ bp) | 45 | **4** | 44 |
| Total length ($\geq 0$ bp) | 5340498 | 5212355 | **5383836** |
| Total length ($\geq 1000$ bp) | 5340498 | 5212355 | **5383836** |
| # contigs | 45 | **4** | 44 |
| Largest contig | 4097401 | **5168551** | 4099674 |
| Total length | 5340498 | 5212355 | **5383836** |
| Reference length | 5167383 | 5167383 | 5167383 |
| GC (%) | 42.16 | 42.06 | 42.19 |
| Reference GC (%) | 42.05 | 42.05 | 42.05 |
| N50 | 4097401 | **5168551** | 4099674 |
| NG50 | 4097401 | **5168551** | 4099674 |
| N75 | 4097401 | **5168551** | 4099674 |
| NG75 | 4097401 | **5168551** | 4099674 |
| L50 | 1 | 1 | 1 |
| LG50 | 1 | 1 | 1 |
| L75 | 1 | 1 | 1 |
| LG75 | 1 | 1 | 1 |
| # misassemblies | **1** | **1** | 3 |
| # misassembled contigs | **1** | **1** | 2 |
| Misassembled contigs length | **9679** | **9679** | 4117533 |
| # local misassemblies | **2** | 3 | 4 |
| # unaligned contigs | 39 + 0 part | **1 + 0 part** | 39 + 0 part |
| Unaligned length | 135514 | **17453** | 163702 |
| Genome fraction (%) | 100.000 | 100.000 | 100.000 |
| Duplication ratio | 1.007 | **1.005** | 1.010 |
| # N's per 100 kbp | 4.25 | **0.48** | 4.46 |
| # mismatches per 100 kbp | 9.56 | **1.30** | 10.14 |
| # indels per 100 kbp | 92.62 | **54.90** | 94.75 |
| Largest alignment | 4097400 | **5168480** | 4098915 |
| NA50 | 4097400 | **5168480** | 4098915 |
| NGA50 | 4097400 | **5168480** | 4098915 |
| NA75 | 4097400 | **5168480** | 4098915 |
| NGA75 | 4097400 | **5168480** | 4098915 |
| LA50 | 1 | 1 | 1 |
| LGA50 | 1 | 1 | 1 |
| LA75 | 1 | 1 | 1 |
| LGA75 | 1 | 1 | 1 |

**Table 4.** (b) in Table 2. All statistics are based on contigs of size≥500 bp, unless otherwise noted (e.g., ”# contigs (≥ 0 bp)” and ”Total length (≥ 0 bp)” include all contigs).

| Assembly | HGAP | FinisherSC | PBJelly |
| --- | --- | --- | --- |
| # contigs ($\geq 0$ bp) | 163 | **41** | 115 |
| # contigs ($\geq 1000$ bp) | 163 | **41** | 115 |
| Total length ($\geq 0$ bp) | 5536634 | 5139491 | **5821106** |
| Total length ($\geq 1000$ bp) | 5536634 | 5139491 | **5821106** |
| # contigs | 163 | **41** | 115 |
| Largest contig | 254277 | **637485** | 495596 |
| Total length | 5536634 | 5139491 | **5821106** |
| Reference length | 5167383 | 5167383 | 5167383 |
| GC (%) | 41.98 | 41.96 | 42.01 |
| Reference GC (%) | 42.05 | 42.05 | 42.05 |
| N50 | 89239 | **215810** | 145441 |
| NG50 | 94672 | **215810** | 161517 |
| N75 | 44568 | **117879** | 98297 |
| NG75 | 53723 | **117879** | 116800 |
| L50 | 20 | **9** | 14 |
| LG50 | 18 | **9** | 12 |
| L75 | 42 | **17** | 26 |
| LG75 | 36 | **17** | 21 |
| # misassemblies | **0** | **0** | 12 |
| # misassembled contigs | **0** | **0** | 10 |
| Misassembled contigs length | **0** | **0** | 439591 |
| # local misassemblies | **0** | 3 | 3 |
| # unaligned contigs | 46 + 1 part | **1 + 0 part** | 43 + 22 part |
| Unaligned length | 200727 | **11862** | 302804 |
| Genome fraction (%) | 98.727 | 98.957 | **99.964** |
| Duplication ratio | 1.046 | **1.003** | 1.068 |
| # N's per 100 kbp | 15.64 | **6.17** | 9.50 |
| # mismatches per 100 kbp | **63.00** | 67.78 | 68.92 |
| # indels per 100 kbp | **577.98** | 589.72 | 597.99 |
| Largest alignment | 254274 | **637485** | 495589 |
| NA50 | 89239 | **215810** | 145441 |
| NGA50 | 94672 | **215810** | 161490 |
| NA75 | 44567 | **117879** | 98293 |
| NGA75 | 50860 | **117879** | 115834 |
| LA50 | 20 | **9** | 14 |
| LGA50 | 18 | **9** | 12 |
| LA75 | 42 | **17** | 26 |
| LGA75 | 36 | **17** | 21 |

**Table 5.** (c) in Table 2. All statistics are based on contigs of size ≥ 500 bp, unless otherwise noted (e.g., ”# contigs (≥ 0 bp)” and ”Total length (≥ 0 bp)” include all contigs).

| Assembly | HGAP | FinisherSC | PBJelly |
| --- | --- | --- | --- |
| # contigs ($\geq 0$ bp) | 21 | **7** | 14 |
| # contigs ($\geq 1000$ bp) | 21 | **7** | 14 |
| Total length ($\geq 0$ bp) | 4689701 | 4660679 | **4718818** |
| Total length ($\geq 1000$ bp) | 4689701 | 4660679 | **4718818** |
| # contigs | 21 | **7** | 14 |
| Largest contig | 1241016 | **2044060** | 1958341 |
| Total length | 4689701 | 4660679 | **4718818** |
| Reference length | 4639221 | 4639221 | 4639221 |
| GC (%) | 50.87 | 50.85 | 50.85 |
| Reference GC (%) | 50.79 | 50.79 | 50.79 |
| N50 | 392114 | **1525398** | 1200847 |
| NG50 | 392114 | **1525398** | 1200847 |
| N75 | 252384 | **1525398** | 275618 |
| NG75 | 252384 | **1525398** | 321636 |
| L50 | 3 | **2** | **2** |
| LG50 | 3 | **2** | **2** |
| L75 | 7 | **2** | 4 |
| LG75 | 7 | **2** | 3 |
| # misassemblies | **8** | **8** | 12 |
| # misassembled contigs | 4 | **3** | 5 |
| Misassembled contigs length | **2530799** | 3584781 | 3672462 |
| # local misassemblies | **3** | **3** | 4 |
| # unaligned contigs | 0 + 1 part | **0 + 0 part** | 0 + 3 part |
| Unaligned length | 205 | **0** | 2605 |
| Genome fraction (%) | 99.583 | 99.656 | **99.689** |
| Duplication ratio | 1.015 | **1.008** | 1.021 |
| # N's per 100 kbp | 0.00 | 0.00 | 0.00 |
| # mismatches per 100 kbp | **3.66** | 4.61 | 4.11 |
| # indels per 100 kbp | **8.36** | 13.82 | 10.75 |
| Largest alignment | 683967 | **1094192** | 949307 |
| NA50 | 339478 | **860437** | 685586 |
| NGA50 | 339478 | **860437** | 685586 |
| NA75 | 229039 | **378942** | 255377 |
| NGA75 | 229039 | **378942** | 255377 |
| LA50 | 5 | **3** | **3** |
| LGA50 | 5 | **3** | **3** |
| LA75 | 9 | **5** | 7 |
| LGA75 | 9 | **5** | 7 |

**Table 6.** (d) in Table 2. All statistics are based on contigs of size≥ 500 bp, unless otherwise noted (e.g., ”# contigs (≥ 0 bp)” and ”Total length (≥ 0 bp)” include all contigs).

| Assembly | HGAP | FinisherSC | PBJelly |
| --- | --- | --- | --- |
| # contigs ($\geq 0$ bp) | 3 | **1** | 2 |
| # contigs ($\geq 1000$ bp) | 3 | **1** | 2 |
| Total length ($\geq 0$ bp) | 3102769 | 3099349 | **3106774** |
| Total length ($\geq 1000$ bp) | 3102769 | 3099349 | **3106774** |
| # contigs | 3 | **1** | 2 |
| Largest contig | 1390744 | **3099349** | 1715191 |
| Total length | 3102769 | 3099349 | **3106774** |
| Reference length | 3097457 | 3097457 | 3097457 |
| GC (%) | 63.38 | 63.39 | 63.38 |
| Reference GC (%) | 63.38 | 63.38 | 63.38 |
| N50 | 1053479 | **3099349** | 1715191 |
| NG50 | 1053479 | **3099349** | 1715191 |
| N75 | 1053479 | **3099349** | 1391583 |
| NG75 | 1053479 | **3099349** | 1391583 |
| L50 | 2 | **1** | **1** |
| LG50 | 2 | **1** | **1** |
| L75 | 2 | **1** | 2 |
| LG75 | 2 | **1** | 2 |
| # misassemblies | 0 | 0 | 0 |
| # misassembled contigs | 0 | 0 | 0 |
| Misassembled contigs length | 0 | 0 | 0 |
| # local misassemblies | 2 | 2 | 2 |
| # unaligned contigs | 0 + 0 part | 0 + 0 part | 0 + 0 part |
| Unaligned length | 0 | 0 | 0 |
| Genome fraction (%) | 99.966 | **99.986** | **99.986** |
| Duplication ratio | 1.002 | **1.001** | 1.003 |
| # N's per 100 kbp | 0.00 | 0.00 | 0.00 |
| # mismatches per 100 kbp | **0.03** | 0.26 | 0.10 |
| # indels per 100 kbp | **1.10** | 3.87 | 1.32 |
| Largest alignment | 1390744 | **3099004** | 1713236 |
| NA50 | 1053134 | **3099004** | 1713236 |
| NGA50 | 1053134 | **3099004** | 1713236 |
| NA75 | 1053134 | **3099004** | 1391558 |
| NGA75 | 1053134 | **3099004** | 1391558 |
| LA50 | 2 | **1** | **1** |
| LGA50 | 2 | **1** | **1** |
| LA75 | 2 | **1** | 2 |
| LGA75 | 2 | **1** | 2 |

**Table 7.** (e) in Table 2. All statistics are based on contigs of size≥ 500 bp, unless otherwise noted (e.g., ”# contigs (≥ 0 bp)” and ”Total length (≥ 0 bp)” include all contigs).

| Assembly | HGAP | FinisherSC | PBJelly |
| --- | --- | --- | --- |
| # contigs ($\geq 0$ bp) | 18 | **5** | 8 |
| # contigs ($\geq 1000$ bp) | 18 | **5** | 8 |
| Total length ($\geq 0$ bp) | 5184825 | 5167414 | **5210862** |
| Total length ($\geq 1000$ bp) | 5184825 | 5167414 | **5210862** |
| # contigs | 18 | **5** | 8 |
| Largest contig | 2103385 | 2913716 | **3343452** |
| Total length | 5184825 | 5167414 | **5210862** |
| Reference length | 5167383 | 5167383 | 5167383 |
| GC (%) | 42.05 | 42.05 | 42.07 |
| Reference GC (%) | 42.05 | 42.05 | 42.05 |
| N50 | 1403814 | 2913716 | **3343452** |
| NG50 | 1403814 | 2913716 | **3343452** |
| N75 | 790287 | **2225895** | 1814491 |
| NG75 | 790287 | **2225895** | 1814491 |
| L50 | 2 | **1** | **1** |
| LG50 | 2 | **1** | **1** |
| L75 | 3 | **2** | **2** |
| LG75 | 3 | **2** | **2** |
| # misassemblies | **1** | **1** | 2 |
| # misassembled contigs | **1** | **1** | 2 |
| Misassembled contigs length | **1403814** | 2913716 | 1820739 |
| # local misassemblies | **0** | **0** | 1 |
| # unaligned contigs | **0 + 0 part** | **0 + 0 part** | 0 + 3 part |
| Unaligned length | **0** | **0** | 13698 |
| Genome fraction (%) | 99.900 | 99.934 | **99.954** |
| Duplication ratio | 1.005 | **1.001** | 1.007 |
| # N's per 100 kbp | 0.00 | 0.00 | 0.00 |
| # mismatches per 100 kbp | 3.41 | 3.56 | **2.28** |
| # indels per 100 kbp | **2.91** | 5.31 | 5.21 |
| Largest alignment | 2103385 | 2225895 | **3343452** |
| NA50 | 1259090 | 1656831 | **3343452** |
| NGA50 | 1259090 | 1656831 | **3343452** |
| NA75 | 790287 | **1656831** | 1270970 |
| NGA75 | 790287 | **1656831** | 1270970 |
| LA50 | 2 | 2 | **1** |
| LGA50 | 2 | 2 | **1** |
| LA75 | 3 | **2** | **2** |
| LGA75 | 3 | **2** | **2** |

### 4.2 Typical use cases

In this section, we describe example use cases of FinisherSC. Below are several scenarios that FinisherSC is helpful to you.

#### 4.2.1 Low coverage data

There are many reasons that you end up having low coverage reads. You may want to save chemicals, the genome may be too long, some parts of the experimental setup may just malfunction or you do not want to overwhelm the assembler with huge amount of data. In any of these situations, you want to utilize as much information from the reads as possible because of the scarcity of read data.

#### 4.2.2 Simple setup for assemblers

There are normally a lot of parameters that can be tuned for modern assemblers. It is also often not clear what parameters work best for your data. However, you do not want to waste time in repeatedly running the assembler by varying different combinations of parameters/setting. In this case, you need a tool that can efficiently and automatically improve your assemblies from the raw reads without rerunning the assembler.

#### 4.2.3 Scaffolding

You may have long contigs prepared from one library and long reads prepared from the other. In this case, you want to robustly and seamlessly combine data from two libraries through scaffolding.

### 4.3 Instructions on using FinisherSC

Our software, FinisherSC, is helpful for the use cases discussed above. It processes long contigs with long reads. You only need to supply the input data files and issue a one-line command as follows to perform the processing. Let us assume that mumP is the path to your MUMMER and destP is the location where the input and output files stay.

- Input : raw_reads.fasta, contigs.fasta
- Output : improved3.fasta
- Command :

~~~
python finisherSC.py destP/ mumP/
~~~

We provide a sandbox example linked in our webpage. Besides the standard usage, there are extra options with details in our webpage.Specifically, we note that users can run FinisherSC in parallel by using the option of [-par numberOfThreads].

### 4.4 Detailed description of the algorithm

We adopt the terminology in Lam et al. (2014a). Random flanking region refers to the neighborhood of a repeat interior. A copy of a repeat being bridged means that some reads cover the copy into the random flanking region. Subroutine 1 removes embedded contigs that would otherwise confuse the later string graph operations. Subroutines 2, 3, 6, 7 are designed to handle repeats. Subroutines 2, 3 resolve repeats whose copies are all bridged by some reads. Subroutines 6, 7 resolve two-copies repeats of which only one copy is bridged. Subroutines 4, 5 utilize neglected information from raw reads. They define merges at locations which are not parts of any long repeats. ^1^

#### Algorithm 2: Subroutine 1: Filter completely embedded contigs

**Input** : contigs.fasta

**Output** : noEmbed.fasta

1. Obtain alignment among contigs from contigs.fasta
2. For any (x,y) contig pair, if x is completely embedded in y, then we add x to removeList
3. Remove all contigs specified in removetList from contigs.fasta. The resultant set of contigs are outputted as noEmbed.fasta

#### Algorithm 3: Subroutine 2: Form a string graph with the BEST successors/predecessors as edges

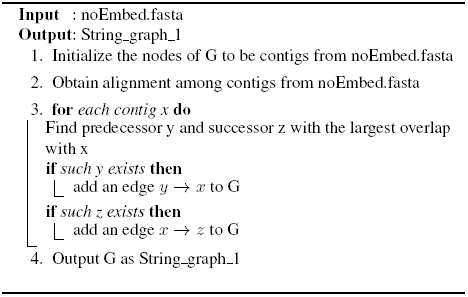

#### Algorithm 4: Subroutine 3: Condense the string graph by contracting edges with both in-degree and out-degree being 1

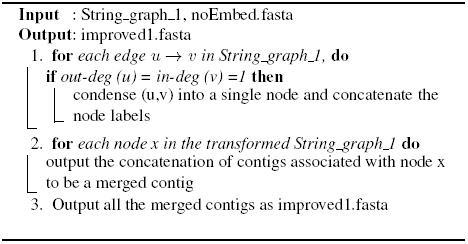

#### Algorithm 5: Subroutine 4: Use raw reads to declare potential successors/predecessors of dangling contigs

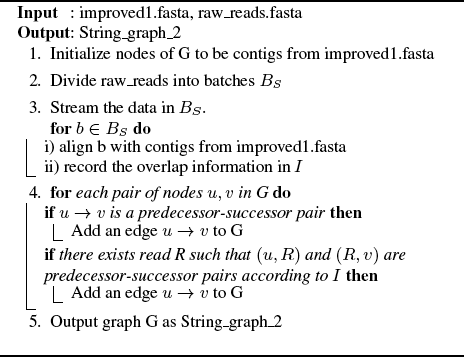

### 4.5 Justification of the algorithm

#### 4.5.1 Big picture

There are two main parts of the algorithm underlying FinisherSC. They are

1. Gap filling
2. Repeat resolution

With uniform sampling assumption, the gaps are unlikely to land on the few long repeats on bacterial genomes. Therefore, subroutines 4, 5 can close most of the gaps. For repeat resolution, subroutines 1, 2, 3, 6, 7 robustly define merges using various transformations of string graphs. Detailed discussion is in the coming section.

##### Algorithm 6: Subroutine 5: Merge contigs (with gaps filled by reads) when they respectively only have 1 successor/1 predecessor

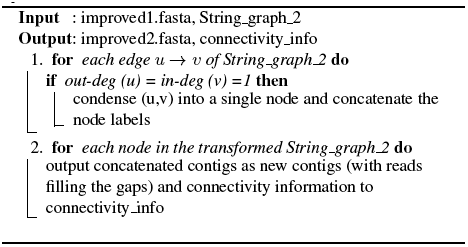

##### Algorithm 7: Subroutine 6: Form a string graph with ALL successors/predecessors as edges

**Input** : improved2.fasta, connectivity_info

**Output** : String_graph_3

1. Use connectivity info to form a graph G with nodes from improved2.fasta. All predecessor-successor pairs are edges in G.
2. Output the corresponding graph as String_graph_3

#### 4.5.2 Detailed justification on repeat resolution

We focus the discussion on a long repeat with two copies. To simplify discussion, we further assume that each base of the genome is covered by some reads and the read length is fixed. The goal here is to correctly merge as many reads as possible in the presence of that repeat. The claim is that Subroutines 2, 3, 6, 7 collectively can achieve this goal. In the case of one repeat, we only need consider the reads either bridging the repeat copies/ reads at the interior of repeats/ touching the repeat copies of that repeat. We separate the discussion on each of the cases depicted in the rows of Fig 4. They are listed as follows.

1. Both copies are bridged
2. Only one copy is bridged
3. Both copies are not bridged

**Fig. 4.**
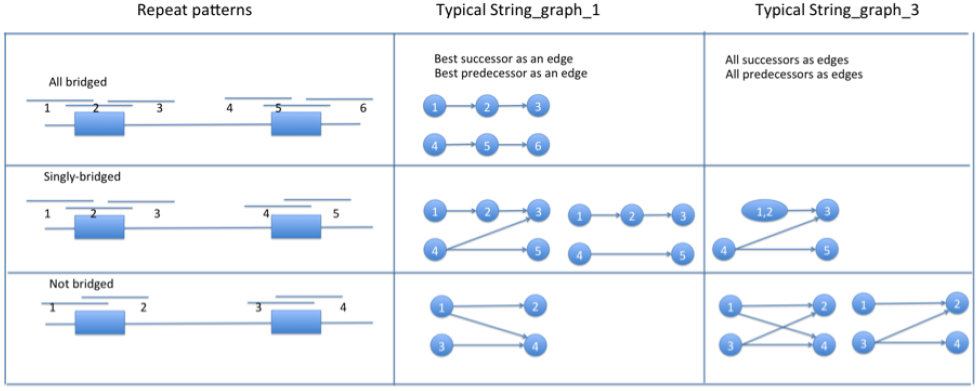
Repeat patterns, typical String_graph_1, typical String_graph_3

##### Algorithm 8: Subroutine 7: Merge contigs with only 1 predecessor or 1 successor and each has no more than two competing edges

**Input** : improved2.fasta, String_graph_3

**Output** :improved3.fasta

1. Traverse the String_graph-3 for pattern of *u*1 → *u*3, *u*2 → *u*3, *u*2 → *u*4 and that out-deg (u1) = 1, out-deg (u2) = 2, in-deg (u3) = 2, in-deg (u4) = 1, if found, then,

a. Delete the edge *u*2 → *u*3
b. Condense the graph
c. Continue until the whole graph is traversed
2. Output the merged contigs as improved3.fasta

Let us first clarify some terminologies before proceeding. A read y is called a successor of another read x if they have significant head-to-tail overlap in the order of *x→y*. y is called the best successor of x if the overlap length is the largest among all the successors of x. y is called the true immediate successor of x if y is the closest read to x’s right side in the sequencing process. Similarly, we can also define the corresponding notion for predecessors.

In the first case, without loss of generality, let us consider any read R emerging from the left flanking region of the left copy. It will get merged with its best successor when condensing String_graph_1. Moreover, the best successor is also the true immediate successor. It is because reads from the other copy of the repeat either have smaller overlap length or are not successors.

Now, let us move to the second case. Since there is a bridging read, there are no reads completely embedded in the interior of the repeat. Without loss of generality, we consider the case that the left copy is bridged and the right copy is not. Now we label R2 as the bridging read, R1/R3 respectively as the true immediate predecessor/successor of the bridging read, R4/R5 as the most penetrating reads into the second copy of the repeat. For all other reads, they get merged with their true immediate successors/predecessors when condensing in String_graph_1. For the remaining five items of interest, the main question is whether there is an edge between R4 and R5 in String_graph_1 (i.e. whether the best successor of R4 is R3). If not, then condensing in String_graph_1 will merge R4 with R5, which is the true immediate successor. If such an edge exists, then we end up with the pattern shown in Fig 4 for String_graph_3. This means that only R1 is merged to R2 when condensing String_graph_1. However, given the existence of the Z-shape pattern, graph operations on String_graph_3, the subroutine 7 will merge R2 and R3, and also will merge R4 and R5.

Finally, consider the third case, when both repeat copies are not bridged. For reads that are not closest to the repeat copies, they get merged correctly when condensing String_graph_1. Without loss of generality, we consider a read x closest to the left flanking region of the left copy of the repeat. An illustration of this situation in String_graph_1 is shown in Fig 5. Let its true immediate successor be T. We are going to show that it will not get merged with the wrong read in String_graph_1 through a proof by contradiction. If x got merged with some wrong F, then *x* → *F* would be an edge. Let y be the read closest the left flanking region of the right copy of the repeat. Then, *y* → *F* is also an edge. Therefore, there should be no merges of *x* → *F*, which results in contradiction. Now we consider String_graph_3, if x has only 1 successor, then it should be T. Otherwise, it is connected to both T and some F. Then, we consider the y coming from the left flanking region of the right copy. There must be an edge from y to F. If there is also an edge from y to T, then both x and y are not merged in String_graph_3. However, if there is no edge from y to T, then x is merged with T and y with F correctly.

**Fig. 5.**
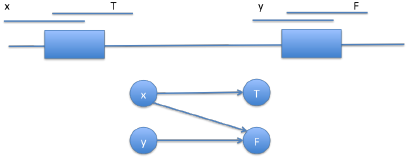
Z-pattern in string graph).

### 4.6 Approximate repeat phasing option

FinisherSC provides an optional component, X-phaser to resolve approximate repeat with two copies, which cannot be bridged by any reads. An example of such an approximate repeat is shown in Fig 6. The algorithm behind X-phaser involves two main parts.

**Fig. 6.**
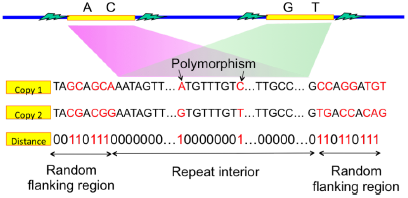
Approximate repeat with two copies

1. Identify repeat interior and its flanking regions
2. Merge contigs by phasing the polymorphisms within the repeat

Algorithm 9 achieves the first part by performing various operations on a string graph. The nodes of the string graph are either contigs or reads near the end points of the contigs. An illustration of a typical string graph is shown in Fig 7. The contigs are indicated by solid circles and reads are indicated by rectangles. The dotted circles specify the random flanking region and repeat interior that we want to infer through Algorithm 9. The X-phasing step in Lam et al. (2014a) achieves the second part. It utilizes the polymorphisms within the repeat interior to help distinguish the repeat copies. Interested readers may refer to Xphased-Multibridging in Lam et al. (2014a) for more details. In FinisherSC, we use the implementation of the Xphasing step in Lam et al. (2014a).

**Fig. 7.**
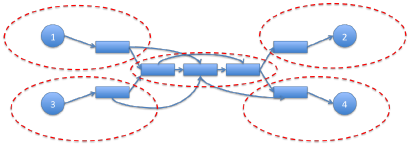
Using string graph to define repeat interiors and flanking regions

### 4.7 Tandem repeat resolution option

The optional tandem repeat resolution step can resolve back-to-back tandem repeat. A back-to-back tandem repeat refers to repeat whose copies directly follow one another. An example is given in Fig 8. FinisherSC provides a component, T-solver, to resolve that. The key idea is to detect cycles in the string graph that join the reads and contigs. An illustration of such a graph is shown in Fig 9, where the circles are contigs and rectangles are reads associated with the end points of the contigs. The algorithm behind T-solver is in Alg 10.

**Fig. 8.**
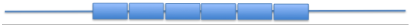
Pattern of back-to-back tandem repeat

**Fig. 9.**
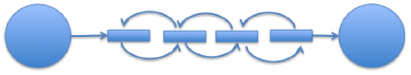
Z-pattern in string graph).

### 4.8 Detailed experimental results on bacterial genomes

In this section, we provide the detailed Quast report for the results described in Table 2. Moreover, we compare in Fig 10 the memory consumption and running time of FinisherSC with those of PBJelly. The computing experiments for this section were performed on the genepool cluster at JGI. Below are commands used to run PBJelly.

~~~
Jelly.py setup Protocol.xml -x “--minGap=1”
~~~

~~~
Jelly.py mapping Protocol.xml
~~~

~~~
Jelly.py support Protocol.xml -x “ --debug”
~~~

~~~
Jelly.py extraction Protocol.xml
~~~

~~~
Jelly.py assembly Protocol.xml -x “--nproc=16“
~~~

**Fig. 10.**
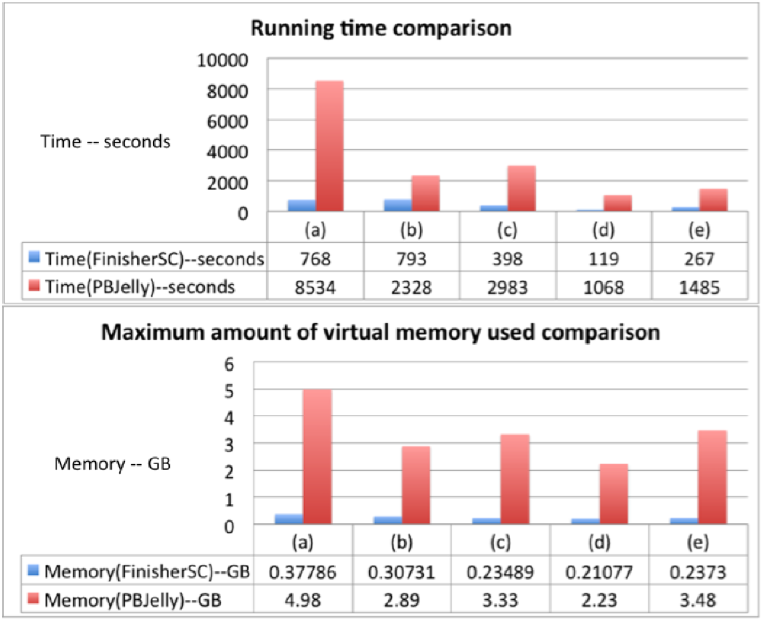
Running time and memory consumption comparison of FinisherSC and PBJelly. (a) to (e) are the corresponding data sets in Table 2.

#### Algorithm 9: Repeat phasing option

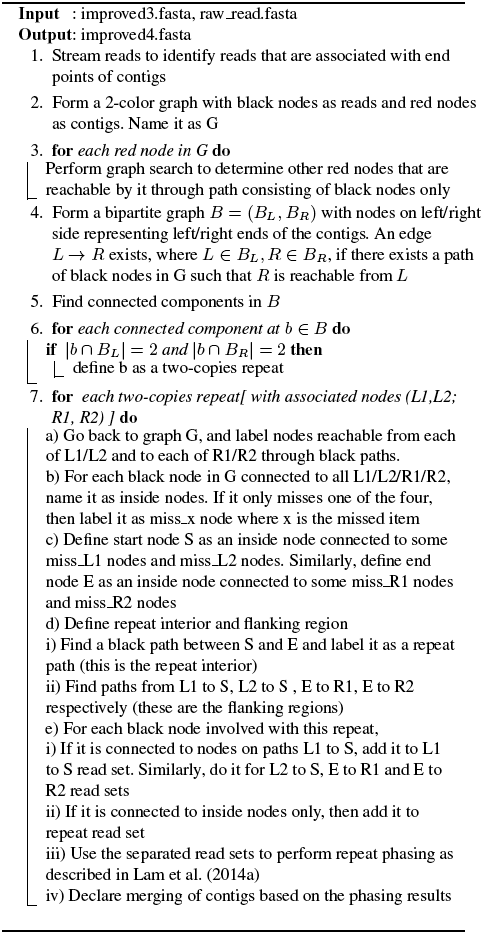

~~~
Jelly.py output Protocol.xml
~~~

The BLASR configuration in Protocol.xml is

~~~
–minMatch 8 –minPctIdentity 70 -bestn 8
~~~

~~~
–nCandidates 30 –maxScore –500 –nproc 16
~~~

~~~
–noSplitSubreads
~~~

### 4.9 Details of the scalability experiments

We run the scalability experiments on a server computer, which is equipped with 64 cores of CPU at clock rate of 2.4-3.3GHz and 512GB of RAM. We also note that, for even larger contig or read data with genomes of higher repeat content, one may be interested in the following options. They are [-f True] for fast alignment and [-l True] for breaking down large contig file. As a reference, we also attach the Quast analysis results on all the intermediate output for the scalability test in Table 8, Table 9 and Table 10. We note that the misassembly count in the Quast analysis for these genomes should only be used as a reference because there is a lack of high quality reference and reference genomes may be from different strains.

**Table 8.** Quast analysis report of Caenorhabditis elegans

| Assembly | contigs | noEmbed | improved | improved2 | improved3 |
| --- | --- | --- | --- | --- | --- |
| # contigs ($\geq 0$ bp) | 245 | 237 | 207 | 200 | **196** |
| # contigs ($\geq 1000$ bp) | 245 | 237 | 207 | 200 | **196** |
| Total length ($\geq 0$ bp) | **104169699** | 103975584 | 103804734 | 103826835 | 103822380 |
| Total length ($\geq 1000$ bp) | **104169699** | 103975584 | 103804734 | 103826835 | 103822380 |
| # contigs | 245 | 237 | 207 | 200 | **196** |
| Largest contig | 3165643 | 3165643 | **4217234** | **4217234** | **4217234** |
| Total length | **104169699** | 103975584 | 103804734 | 103826835 | 103822380 |
| Reference length | 100286401 | 100286401 | 100286401 | 100286401 | 100286401 |
| GC (%) | 35.67 | 35.66 | 35.66 | 35.66 | 35.66 |
| Reference GC (%) | 35.44 | 35.44 | 35.44 | 35.44 | 35.44 |
| N50 | 1613475 | 1613475 | 1773224 | **1832604** | **1832604** |
| NG50 | 1619480 | 1619480 | 1832604 | **1910174** | **1910174** |
| N75 | 834223 | 834223 | 947068 | **959597** | **959597** |
| NG75 | 881109 | 881109 | 959597 | **1031901** | **1031901** |
| L50 | 24 | 24 | 22 | **21** | **21** |
| LG50 | 23 | 23 | 21 | **20** | **20** |
| L75 | 48 | 48 | 43 | **41** | **41** |
| LG75 | 44 | 44 | 40 | **38** | **38** |
| # misassemblies | 1358 | **1334** | 1411 | 1407 | 1403 |
| # misassembled contigs | 127 | 120 | 108 | 104 | **102** |
| Misassembled contigs length | 96744440 | **96571278** | 98809318 | 99454743 | 99479049 |
| # local misassemblies | 784 | **783** | 809 | 805 | 810 |
| # unaligned contigs | 70 + 20 part | 70 + 20 part | 70 + 11 part | 68 + 11 part | **68 + 10 part** |
| Unaligned length | 1862487 | 1862487 | 1743676 | 1745585 | **1743404** |
| Genome fraction (%) | 99.525 | 99.525 | 99.531 | **99.535** | **99.535** |
| Duplication ratio | 1.027 | 1.025 | **1.024** | **1.024** | **1.024** |
| # N's per 100 kbp | 0.00 | 0.00 | 0.00 | 0.00 | 0.00 |
| # mismatches per 100 kbp | 13.27 | 13.27 | 13.23 | 13.20 | **13.13** |
| # indels per 100 kbp | 20.89 | 20.89 | **20.63** | 20.80 | 20.86 |
| Largest alignment | 1362562 | 1362562 | 1362562 | 1362562 | 1362562 |
| NA50 | 407833 | 407833 | **407888** | 402635 | 402635 |
| NGA50 | **421272** | **421272** | **421272** | 412370 | 412370 |
| NA75 | 213728 | **217528** | 213728 | 211961 | 211961 |
| NGA75 | **232043** | **232043** | 229862 | 226413 | 226413 |
| LA50 | 85 | 85 | 85 | 85 | 85 |
| LGA50 | 80 | 80 | 80 | 80 | 80 |
| LA75 | 169 | **168** | **168** | 169 | 169 |
| LGA75 | **156** | **156** | **156** | 157 | 157 |

**Table 9.** Quast analysis report of Drosophila

| Assembly | contigs | noEmbed | improved | improved2 | improved3 |
| --- | --- | --- | --- | --- | --- |
| # contigs ($\geq 0$ bp) | 128 | 128 | 110 | 96 | **93** |
| # contigs ($\geq 1000$ bp) | 128 | 128 | 110 | 96 | **93** |
| Total length ($\geq 0$ bp) | **138490501** | **138490501** | 138082066 | 138131721 | 138096303 |
| Total length ($\geq 1000$ bp) | **138490501** | **138490501** | 138082066 | 138131721 | 138096303 |
| # contigs | 128 | 128 | 110 | 96 | **93** |
| Largest contig | 24648237 | 24648237 | **27967410** | **27967410** | **27967410** |
| Total length | **138490501** | **138490501** | 138082066 | 138131721 | 138096303 |
| Reference length | 168717020 | 168717020 | 168717020 | 168717020 | 168717020 |
| GC (%) | 41.86 | 41.86 | 41.87 | 41.87 | 41.87 |
| Reference GC (%) | 41.74 | 41.74 | 41.74 | 41.74 | 41.74 |
| N50 | 15305620 | 15305620 | **21710673** | **21710673** | **21710673** |
| NG50 | 6168915 | 6168915 | **15305620** | **15305620** | **15305620** |
| N75 | 920983 | 920983 | 1012145 | **1448033** | **1448033** |
| NG75 | 291391 | 291391 | 308285 | 389196 | **402690** |
| L50 | 4 | 4 | **3** | **3** | **3** |
| LG50 | 6 | 6 | **4** | **4** | **4** |
| L75 | 15 | 15 | 11 | **10** | **10** |
| LG75 | 57 | 57 | 50 | 42 | **41** |
| # misassemblies | 8272 | 8272 | **8057** | 8071 | 8064 |
| # misassembled contigs | 98 | 98 | 84 | 76 | **73** |
| Misassembled contigs length | **127872011** | **127872011** | 130423069 | 130352009 | 130316591 |
| # local misassemblies | 290 | 290 | **282** | 283 | 284 |
| # unaligned contigs | 1 + 0 part | 1 + 0 part | 1 + 0 part | 1 + 0 part | 1 + 0 part |
| Unaligned length | 61383 | 61383 | 61383 | 61383 | 61383 |
| Genome fraction (%) | **76.376** | **76.376** | 76.348 | 76.367 | 76.364 |
| Duplication ratio | 1.082 | 1.082 | **1.079** | **1.079** | **1.079** |
| # N's per 100 kbp | 0.00 | 0.00 | 0.00 | 0.00 | 0.00 |
| # mismatches per 100 kbp | 21.85 | 21.85 | **21.34** | 21.81 | 21.84 |
| # indels per 100 kbp | 20.88 | 20.88 | **20.81** | 20.94 | 20.99 |
| Largest alignment | 6494798 | 6494798 | 6494798 | 6494798 | 6494798 |
| NA50 | 1316520 | 1316520 | 1316520 | **1377072** | **1377072** |
| NGA50 | 820949 | 820949 | 821097 | **829069** | **829069** |
| NA75 | 308622 | 308622 | 341648 | **398564** | **398564** |
| NGA75 | **6441** | **6441** | 6389 | 6414 | 6384 |
| LA50 | 26 | 26 | 26 | 26 | 26 |
| LGA50 | 41 | 41 | 41 | **40** | **40** |
| LA75 | 78 | 78 | 76 | **72** | **72** |
| LGA75 | 532 | 532 | 533 | **520** | 522 |

**Table 10.** Quast analysis report of Saccharomyces cerevisiae

| Assembly | contigs | noEmbed | improved | improved2 | improved3 |
| --- | --- | --- | --- | --- | --- |
| # contigs ($\geq 0$ bp) | 30 | 27 | 23 | 22 | **20** |
| # contigs ($\geq 1000$ bp) | 30 | 27 | 23 | 22 | **20** |
| Total length ($\geq 0$ bp) | **12370681** | 12295446 | 12221471 | 12225040 | 12220764 |
| Total length ($\geq 1000$ bp) | **12370681** | 12295446 | 12221471 | 12225040 | 12220764 |
| # contigs | 30 | 27 | 23 | 22 | **20** |
| Largest contig | 1538192 | 1538192 | 1538192 | 1538192 | 1538192 |
| Total length | **12370681** | 12295446 | 12221471 | 12225040 | 12220764 |
| Reference length | 12157105 | 12157105 | 12157105 | 12157105 | 12157105 |
| GC (%) | 38.21 | 38.18 | 38.17 | 38.17 | 38.17 |
| Reference GC (%) | 38.15 | 38.15 | 38.15 | 38.15 | 38.15 |
| N50 | 777787 | 777787 | 777787 | 777787 | 777787 |
| NG50 | 777787 | 777787 | 777787 | 777787 | 777787 |
| N75 | 544615 | 544615 | 544615 | 544615 | **583193** |
| NG75 | 544615 | 544615 | 544615 | 544615 | **583193** |
| L50 | 6 | 6 | 6 | 6 | 6 |
| LG50 | 6 | 6 | 6 | 6 | 6 |
| L75 | 11 | 11 | 11 | 11 | 11 |
| LG75 | 11 | 11 | 11 | 11 | 11 |
| # misassemblies | 112 | 109 | **106** | **106** | 107 |
| # misassembled contigs | 27 | 24 | 20 | 20 | **18** |
| Misassembled contigs length | 11049515 | 10974280 | 10900305 | 10900305 | **10896029** |
| # local misassemblies | 21 | 21 | 21 | 21 | 21 |
| # unaligned contigs | 0 + 0 part | 0 + 0 part | 0 + 0 part | 0 + 0 part | 0 + 0 part |
| Unaligned length | 0 | 0 | 0 | 0 | 0 |
| Genome fraction (%) | 98.117 | 98.117 | 98.215 | **98.243** | **98.243** |
| Duplication ratio | 1.038 | 1.032 | **1.024** | **1.024** | **1.024** |
| # N's per 100 kbp | 0.00 | 0.00 | 0.00 | 0.00 | 0.00 |
| # mismatches per 100 kbp | **76.68** | **76.68** | 78.86 | 79.14 | 78.05 |
| # indels per 100 kbp | 11.71 | 11.71 | **11.67** | 13.58 | 13.50 |
| Largest alignment | 1027016 | 1027016 | 1027016 | **1053518** | **1053518** |
| NA50 | **377112** | **377112** | 377016 | 377016 | 377016 |
| NGA50 | **377112** | **377112** | 377016 | 377016 | 377016 |
| NA75 | 181420 | **198393** | **198393** | **198393** | **198393** |
| NGA75 | 198393 | 198393 | 198393 | 198393 | 198393 |
| LA50 | 11 | 11 | 11 | 11 | 11 |
| LGA50 | 11 | 11 | 11 | 11 | 11 |
| LA75 | 23 | **22** | **22** | **22** | **22** |
| LGA75 | 22 | 22 | 22 | 22 | 22 |

#### ACKNOWLEDGEMENT

The authors K.K.L and D.T. are partially supported by the Center for Science of Information (CSoI), an NSF Science and Technology Center, under grant agreement CCF-0939370. The work conducted by the U.S. Department of Energy Joint Genome Institute, a DOE Office of Science User Facility, is supported under Contract No. DE-AC02-05CH11231.

#### Algorithm 10: Tandem repeat resolution option

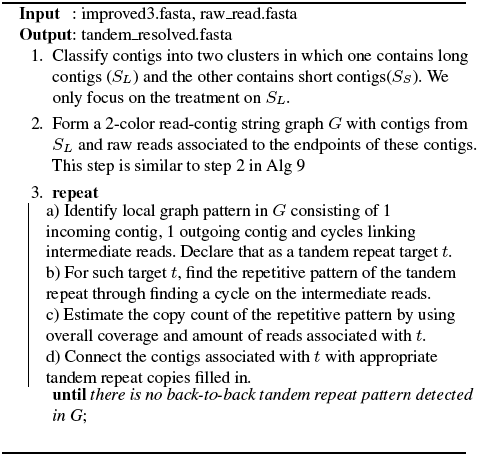

### Footnotes

1 To simplify discussion, the subroutines described are based on the assumption that reads are extracted from a single-stranded DNA. However, we remark that we have implemented FinisherSC by taking into account that reads are extracted from both forward and reverse strands.

## REFERENCES

Chin, C.-S., Alexander, D. H., Marks, P., Klammer, A. A., Drake, J., Heiner, C., Clum, A., Copeland, A., Huddleston, J., Eichler, E. E., et al. (2013). Nonhybrid, finished microbial genome assemblies from long-read smrt sequencing data. Nature methods.

English, A. C., Richards, S., Han, Y., Wang, M., Vee, V., Qu, J., Qin, X., Muzny, D. M., Reid, J. G., Worley, K. C., et al. (2012). Mind the gap: Upgrading genomes with pacific biosciences rs long-read sequencing technology. PLoS ONE, 7: 47768.

Gurevich, A., Saveliev, V., Vyahhi, N., and Tesler, G. (2013). Quast: quality assessment tool for genome assemblies. Bioinformatics, 29(8):1072–1075.

Kurtz, S., Phillippy, A., Delcher, A. L., Smoot, M., Shumway, M., Antonescu, C., and Salzberg, S. L. (2004). Versatile and open software for comparing large genomes. Genome biology, 5(2):R12.

Lam, K.-K., Khalak, A., and Tse, D. (2014a). Near-optimal assembly for shotgun sequencing with noisy reads. BMC Bioinformatics.

Lam, K.-K., LaButti, K., Khalak, A., and Tse, D. (2014b). http://kakitone.github.io/finishingTool/.

Myers, E. W., Sutton, G. G., Delcher, A. L., Dew, I. M., Fasulo, D. P., Flanigan, M. J., Kravitz, S. A., Mobarry, C. M., Reinert, K. H., Remington, K. A., et al. (2000). A whole-genome assembly of drosophila.

